# Improved Microbubble Tracking for Super-Resolution Ultrasound Localization Microscopy using a Bi-Directional Long Short-term Memory Neural Network

**DOI:** 10.1101/2025.02.10.637352

**Authors:** Xi Chen, Matthew R. Lowerison, YiRang Shin, Yike Wang, Zhijie Dong, Qi You, Pengfei Song

## Abstract

Ultrasound localization microscopy (ULM) enabled high-accuracy measurements of microvessel flow beyond the resolution limit of conventional ultrasound imaging by utilizing contrast microbubbles (MBs) as point targets. Robust tracking of MBs is an essential task for fast and high-quality ULM image reconstruction. Existing MB tracking methods suffer from challenging imaging scenarios such as high-density MB distributions, fast blood flow, and complex flow dynamics. Here we present a deep learning-based MB pairing and tracking method based on a bi-directional long short-term memory neural network for ULM. The proposed method integrates multiparametric MB characteristics to facilitate more robust and accurate MB pairing and tracking. The method was validated on a simulation data set, a tissue-mimicking flow phantom, and *in vivo* on a mouse and rat brain.

## I. Introduction

**Q**uantitative characterization of hemodynamics provides powerful biomarkers for studying various physiological functions as well as assessing health conditions. Conventional Doppler ultrasound earned its important role in clinic via the ability to measure blood flow noninvasively and quantitatively in real time [1]. However, existing Doppler blood flow imaging technologies suffer from low spatial resolution due to the diffraction limit of ultrasound as well as measurement errors associated with Doppler angle [2]. Overcoming these limitations remains an important research topic in ultrasound blood flow imaging. Ongoing research efforts have been made to enhance the sensitivity to small vessels, as well as reducing the angle dependency of Doppler velocity measurements [3].

Ultrasound localization microscopy (ULM) [4], [5] is a contrast microbubble (MB)-based super-resolution imaging technique that gained popularity over the past decade [6], [7], [8]. Inspired by the principles of localization microscopy in optical imaging (e.g., PALM [9], [10] and STORM [11]), ULM utilizes MBs as point targets to break the acoustic diffraction limit and achieve super-resolution. MB localization and MB tracking are the two major steps in the ULM processing workflow. Accumulating high-fidelity estimation of detected MB locations from using ultrafast imaging allows reconstruction of the underlying vascular structure beyond the conventional resolution limit (e.g., ~10X improvement) [12]. In addition to the high spatial resolution, ULM also provides accurate flow velocity measurements by tracking the movement of individual MBs across time, which is independent from Doppler angle [4]. The unique capability of imaging deep tissue microvasculature *in vivo* while providing microvessel flow velocity measurements opened new doors for many preclinical and clinical applications for ULM [6].

While high accuracy MB localization lays the foundation for downstream ULM processing, a robust MB pairing and tracking pipeline is also critical for high quality ULM [13]. The ability to track the same MB consistently and accurately for a sustained period of time ensures reconstruction of ULM microvessel density and flow velocity maps with high confidence. In addition, erroneous MB localizations can also be corrected for or removed by applying appropriate constraints to regulate the MB tracking results [14]. An enhanced MB tracking performance under challenging imaging conditions such as high MB concentration (or high degree of MB signal overlap), fast blood flow, and complex flow dynamics also translates to more efficient use of the acquired MB signals, some of which would have been deemed unreliable and subsequently discarded. In practice, a more efficient utilization of MB signals leads to shorter data acquisition for ULM and improved temporal resolution, which is currently one of the major concerns of ULM in many preclinical and clinical applications.

MB tracking for ULM usually starts with frame-to-frame association of detected MBs, which are then assembled into MB tracks. Early MB tracking algorithms mostly rely on local-window-based searches for each localized MB across consecutive frames [4]. This class of approach involves handling situations where no MB candidate is found within the allowed search radius or choosing the best candidate in regions with dense MB distributions. The search can be solely distance-based [15] or involve correlation coefficient-based template matching between consecutive MB images [5]. The performance of these local-window-based tracking techniques heavily depends on the quality of the MB localization signal, radius selected for the search, as well as any assumptions made on the motion model of the MBs [16], [17]. The computational cost of local-window-based MB tracking approaches also scales with the number of MB localizations and the complexity of the searching conditions [15].

A more simplistic and robust formulation of the MB tracking problem is to use the global pairing approach based on the principles of linear sum assignment (LSA) [13], in which all the MBs between two consecutive imaging frames are paired *globally* by minimizing the total assignment cost (in this case, the cost is the total pair-wise distance between paired MBs). LSA had been thoroughly studied in the fields of optimization and combinatorics [18], with various well-optimized software implementations that are publicly available on different programming platforms [19], [20], [21], [22], [23]. This approach achieves high quality tracking despite its easy implementation and low computational cost. However, existing LSA-based MB tracking approaches mostly rely on spatial distances in pairing, which becomes unstable under high MB concentration or fast blood flow conditions where minimal pairing distance becomes an insufficient condition for reliable MB tracking. Indeed, it was shown in existing study that the accuracy of a pairing-based tracking framework can be greatly improved by integrating additional MB features into the cost function [24]. Nevertheless, the optimal construction of a cost function that incorporates multiple features can vary among different datasets, and the need for handcrafting the combination of features leaves room for human bias. Meanwhile, LSA-based global MB tracking also relies on high frame-rate imaging (e.g., >500 Hz frame rate) to ensure optimal assignment of MB pairs, which may be challenging to realize on most clinical ultrasound imaging systems.

In this paper, we propose a novel solution to enhance the performance of LSA-based global MB frame-to-frame association for robust MB tracking under challenging imaging conditions. Motivated by recent successes of deep learning (DL)-based MB localization [25], [26], [27], [28], [29], [30], [31], we propose a DL-based formulation of the MB pairing and tracking problem utilizing a bi-directional long short-term memory (Bi-LSTM) neural network. The DL-based MB pairing and tracking approach learns to produce an optimal pairing in a data-driven, supervised fashion, reducing empirical biases and uncertainties when designing the rules of pairing. Additionally, we propose to combine the two approaches by using multiple input channels to the Bi-LSTM network with each channel representing a specific MB signal feature to enable a more sophisticated combination of features in the latent space. We systematically tested the performance of each method on simulation data with ground truth, followed by flow phantom experiment and *in vivo* experiment in a mouse brain and a rat brain.

The main contribution of this work is summarized as follows:

- We extended the widely used LSA-based MB pairing by proposing a fusion cost matrix that integrates three features of MB signals: distance, brightness and shape.
- We proposed a DL formulation of the MB pairing problem using a Bi-LSTM neural network that learns to perform MB pairing based on multiple features without the need of manually designing the pairing cost.
- Extensive experimental results on simulation data, flow phantom and *in vivo* mouse and rat brain were presented to show improved robustness and tracking accuracy for both methods. Specifically, our tracking results on mouse and rat brain shows that, despite being trained solely using simulation data, the DL-based method has robust generalizability, outperforming both LSA-based methods (i.e., conventional LSA and LSA with integrated MB signal features).

The rest of the paper is structured as follows: Section II provides a detailed description of proposed methods, as well as our experimental procedure. Section III presents and analyzes our experimental results. Section IV discusses our findings and limitations of the work presented. Section V concludes the paper.

## II. Methods

### A. MB pairing as a linear sum assignment problem

Given two sets of items, the LSA problem finds the minimum total-cost solution of assigning items in the first set to items in the second. It can be formally defined as:

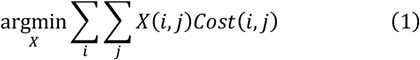

with the constraints ∀*i*: ∑_*j*_ *A*(*i, j*) = 1, ∀*j*: ∑_*i*_ *A*(*i, j*) ≤ 1 if the first set contains more elements, and ∀*i*: ∑ _j_ *A*(*i, j*) = 1, ∀*j*: ∑ _i_ *A*(*i, j*) ≤ 1 otherwise, where *i, j* are indices representing items in the first and the second set, respectively. *X*(*i, j*) is the assignment matrix, where *X*(*i, j*) = 1 if item *i* is assigned to item *j*, and *X*(*i, j*) = 0 otherwise. *Cost*(*i, j*) is the assignment cost matrix, where the (*i, j*)^*th*^ entry represents the cost of assigning item *i* to item *j*.

For the MB pairing problem, the two sets of items are MB detections (i.e., localizations) in two consecutive frames (that is, frame *t* and frame *t*+*1*). Conventional minimum total distance approaches define the cost matrix using the Euclidean distance (for 2D imaging):

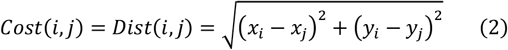

where (*x*_*i*_, *y*_*i*_) is the spatial coordinate of *i*^*th*^ MB localization in frame *t*, and (*x*_*j*_, *y*_*j*_) is the *j*^*th*^ MB spatial coordinate in frame *t*+*1*. This formulation assumes a spatial closeness of the same MB detections in consecutive frames. In practice, when applied to high concentration or fast flowing MB data, the minimization of total pairing distance may no longer provide a satisfactory pairing solution because of the rapid movement and decorrelation of MB signals between consecutive frames. For these challenging cases, our hypothesis is that additional information associated with MB signal characteristics can be utilized to facilitate robust MB pairing performance based on LSA. In addition to the minimum pairing distance, we propose two more assumptions: 1) the amplitude of the MB signal (that is, the amplitude of the backscattered MB signal at the detected MB location) will stay consistent between adjacent consecutive frames and; 2) the shape of the MB signal (i.e., the point-spread-function) will stay consistent within a short period of time. Based on these additional assumptions, the new fusion cost matrix is defined as

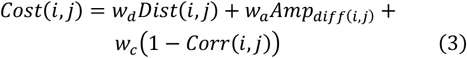

where *Dist*(*i, j*) is the same Euclidean distance as mentioned above, *Amp*_*diff*(*i, j*) is the absolute difference between the amplitude of MB signals at each localized position, and *Corr*(*i, j*) is the normalized correlation coefficient of MB images (within a small patch that contains a single MB) detected at different time points (Fig. 1). All three of the cost terms were normalized to the range of [0, 1] to avoid introducing bias due to the difference in scale. *w*_*d*_, *w*_*a*_ and *w*_*c*_ are weight terms associated with each cost item. We used 1 − *Corr*(*i, j*) so that all lost terms in Eq. (3) are to be minimized. An example of the feature matrices used to construct the fusion cost matrix calculated from consecutive frames of MB detections is shown in **Figure 1**.

**Figure 1.**
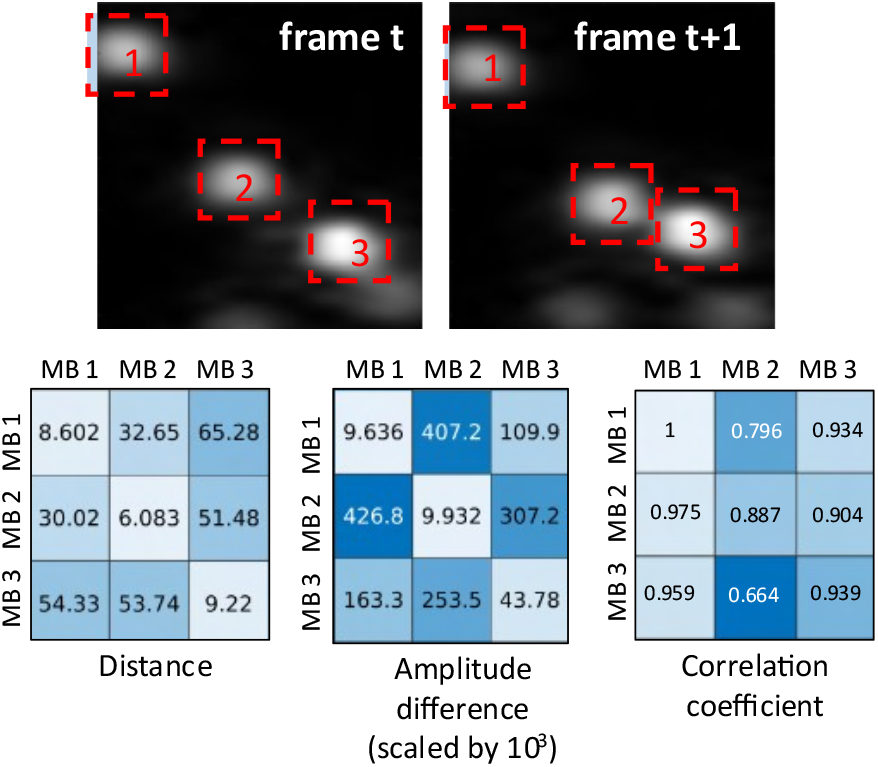
Example of two consecutive frames of simulated MBs with 3 MBs in each frame. Three different features were used to construct the fusion cost matrix: the distance matrix defined as the Euclidean distance between each pair of MB localizations; the amplitude difference matrix defined as the absolute difference of the peak MB signals detected at different time frames (scaled by 10^3^ for display in figure); the normalized correlation coefficient matrix defined as the normalized cross-correlation coefficient based on local patches (red dashed lines) around each MB detection) between each pair of MB detections. The image patches were zero padded (indicated by the shaded area) if desired patches exceed the image boundary.

### B. Bi-LSTM formulation of the linear sum assignment problem

Various studies have demonstrated the feasibility of DL-based formulations of LSA and related data-association problems [32], [33], [34], [35]. Here we followed the setup of the Deep Hungarian Network (DHN) [35] whose input is a pairwise distance matrix and the output approximates the assignment matrix. DHN is well suited for MB pairing and tracking purposes because it can handle input with arbitrary dimensions (ideal for dealing with the varying number of MB detections per frame) and output an assignment matrix with the same dimension as the input. Additionally, we adopted the idea of using Bi-directional LSTM (Bi-LSTM) to extract features necessary to solve the assignment problem with knowledge of the entire input sequence, without having to divide the original problem into sub-assignment problems [32].

Figure 2. shows the architecture of the proposed Bi-LSTM MB pairing network. The input to the network is of spatial dimension M×N, where M is the number of MB detections in frame *t*, and N is the number of detections in frame *t*+*1*. Each pixel value in the matrix represents the cost of assigning one detection in frame *t* and one detection in frame *t*+*1*. Channels of the input matrix correspond to different features used to produce the cost function (Eq. 3). For example, if spatial coordinate is to be used as the channel input, the *ij*^*th*^ element in the channel would be the Euclidian distance between the *i*^*th*^ MB in frame *t* and the *j*^*th*^ MB in frame *t*+*1*. Similar to the LSA methods, values in each channel of the cost matrix were scaled to have a maximum value of 1.

For the data flow in the network, the input is first decomposed into *M* row vectors with a size of *N×3* and subsequently fed into a 1-D Bi-LSTM layer to produce *M* feature vectors of size *N×2H*, where *H* is a parameter associated with the hidden dimensions of LSTM. The channel size increases by a factor of 2 after each Bi-LSTM unit due to concatenation of the forward LSTM output and the backward LSTM output. The feature vectors are concatenated along the column direction to produce the first level hidden representation matrix of size *M×N×2H*.

**Figure 2.**
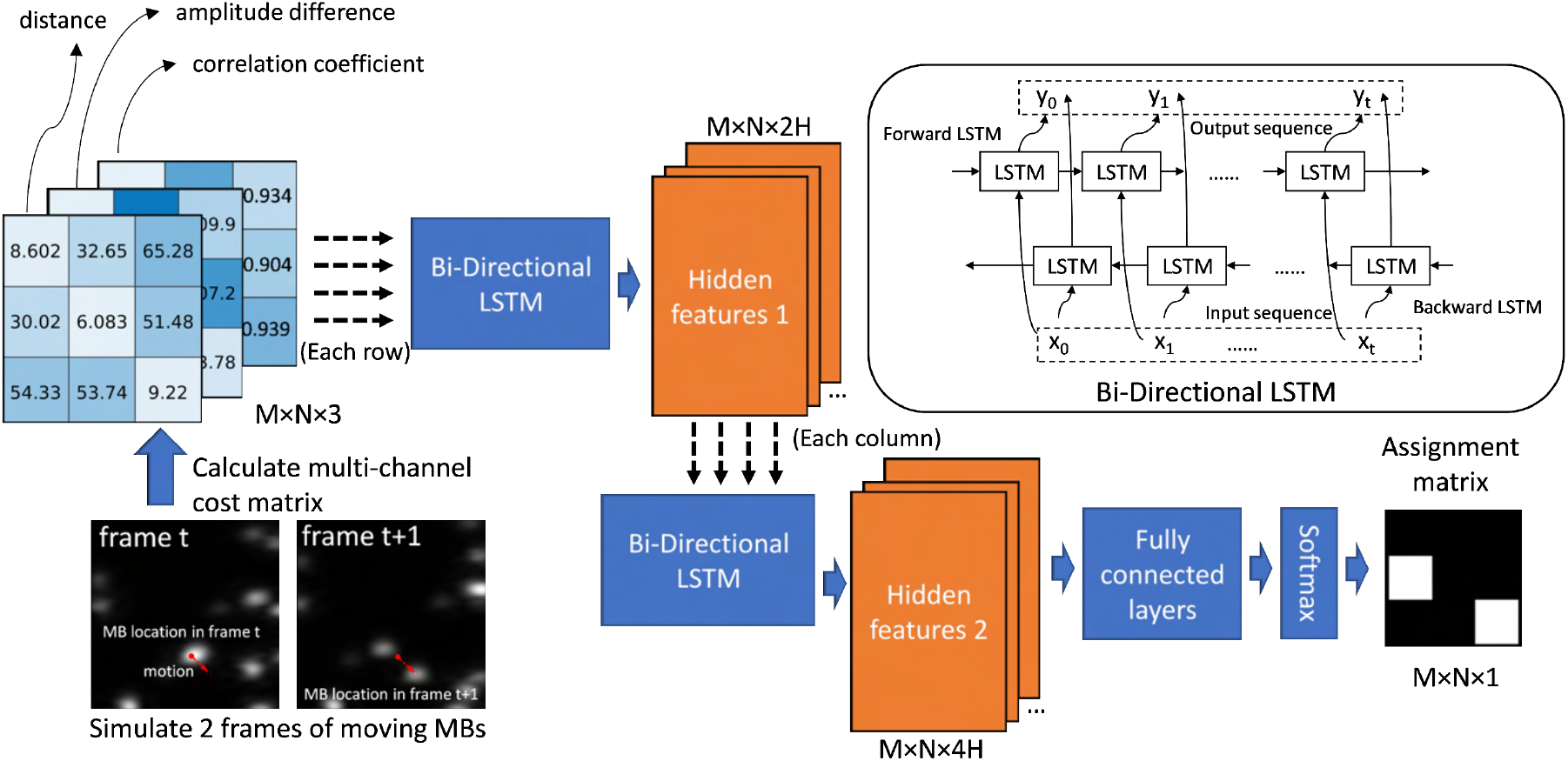
Diagram of the proposed DL formulation of the LSA problem using Bi-LSTM neural networks.

The hidden representation matrix is then broken into *N* column vectors of size *M×2H*. The column vectors are fed into another 1-D Bi-LSTM layer, producing *N* second level features of *M×4H*. The feature vectors are then reshaped and concatenated along the row direction into a second level hidden representation matrix of shape *M×N×4H*. The final part of the network is a sequence of three fully connected (FC) layers operating along the channel dimension, where the final channel size is reduced to one. Softmax was applied to the output of the FC layers to obtain a soft assignment matrix. The design ensures that for any arbitrarily shaped input matrix, the output will be of the same spatial dimension except that the channel dimension will be collapsed.

Compared to vanilla recurrent neural networks (RNNs), LSTM is more capable of forming long-term associations thanks to the use of cell states that act as long-term storage units of information from past events. The Bi-LSTM units can be viewed as feature transformation operations for each element in the cost matrix conditioned on the context of the entire row/column where it resides: for an input of a feature vector of size *n*, at position *i*, the forward LSTM performs inference informed by the cell state, which contains ‘memory’ of input at position 0 through position *i-1*. The cell state is updated recursively through sequential processing of input at position 0 through position *i-1*, as well as the hidden state output of input at position *i-1*. The processing step for *i* will also update the cell state to contain some form of encoding of information associated with input *i*, ensuring that the processing of input at position *i*+*1* is informed by previous input from position 0 to position *i* through the updated cell state. The reverse LSTM processes input in a reversed order: the operation on input at position *i* will be informed by a cell state containing information associated with input at position *n* through position *i*+*1*, as well as the hidden state output of input position *i*+*1*. The combination of outputs from both directions gives us an inference of the input at position *i*, informed by all the input feature vectors excluding position *i*. The row-wise and column-wise operations correspond to a forward pairing search linking MB localizations in frame *t* to those in frame *t*+*1*, and a backward pairing search linking MB localizations from frame *t*+*1* to frame *t*.

### C. Training data generation

In order to produce high-quality training data with realistic ultrasound images of MBs, we used experimentally acquired MB signals to build a point-spread-function (PSF) bank for training data generation. After injecting heavily diluted MB (DEFINITY®, Lantheus Medical Imaging, Inc.) solution into deionized water, ultrasound images of isolated, slow-moving MBs were acquired with a Vantage 256 system (Verasonics Inc., Kirkland, WA) and a L35-16vX high-frequency linear array transducer with a center frequency of 20 MHz and a 9-angle plane wave compounding sequence (1-degree increments). Initial candidates of MB PSF patches were selected by computing the normalized cross-correlation between the MB intensity image frames and an estimated Gaussian PSF and applying a peak detection function ‘imregionalmax.m’. The regions around the extracted peaks were then visually inspected to discard any candidates that are likely to be overlapping MBs or noise. The remaining PSF samples were saved as the PSF bank (**Figure 3**).

**Figure 3.**
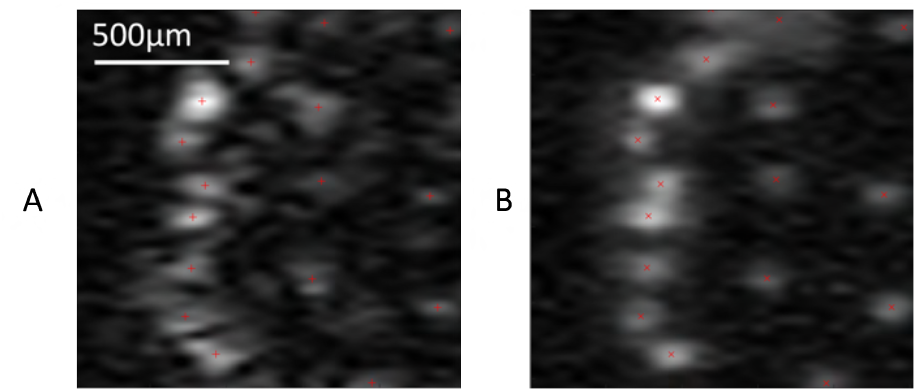
A. *in vivo* ultrasound MB signals collected from a mouse brain. B. Simulated MB signals using experimentally collected MB PSFs.

For simulation, the number of MBs in each frame was randomly drawn from the range of 10-500. A different number of MB localizations between frames was allowed. We then randomly selected a certain number of paired MBs. The initial coordinates of the paired MBs were randomly generated, The MB displacement between frames was restricted to 30 pixels, which corresponds to a displacement of ~1.92 wavelengths in between frames, or ~147mm/s velocity assuming a 4.928µm pixel size and 1000 Hz frame rate. Remaining unpaired MBs were assigned to random locations in both frames. The initial brightness of paired MBs was randomly drawn from a value ranging from [0.2, 1], with a maximum of 20% amplitude fluctuation between frames. The brightness of unpaired MBs was randomly drawn in both frames from the same range of [0.2, 1]. A PSF was randomly drawn from the experimental PSF bank for all the simulated MB signals, with paired MBs having the same PSF in two consecutive frames. The MB image for each individual MB was obtained by convolving the selected PSF patch with an image containing a single point at the desired MB location, scaled by the desired MB brightness. The final MB signals were constructed by adding up all the individual MB images. The ground truth assignment matrix is a matrix comprised of 0s and 1s, with 1s representing paired locations and 0s otherwise. A total of 3000 pairs of consecutive frames were generated for training and 300 pairs for validation.

### D. Neural network training

The neural network was trained by minimizing the focal loss [36], defined as

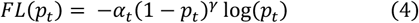

where for an estimated probability of positive class *p* and ground truth label *y, p*_*t*_ = *p* if *y* = 1, and 1 − *p* otherwise. *α* is the term that adjusts the loss to handle class imbalance. *α*_*t*_ = *α* if *y* = 1, and 1 − *α* otherwise. *γ* is the focus parameter. We selected *α* = 0.25, *γ* = 2 empirically.

The network was trained using the Adam optimizer [37] with a learning rate of 0.001 and mini-batch of 10 samples.

### E. MB localization and reconstruction of velocity map

Normalized cross-correlation (NCC) based MB localization was performed to obtain MB locations used for all pairing methods. For each input data acquisition, we estimated a 2-D Gaussian template of the MB PSF based on the appearance of the observed MBs. 2-D NCC template matching was performed using the estimated Gaussian PSF template. MB position estimations were obtained by centroid estimation on the correlation coefficient map.

The reconstruction of the final velocity map begins with formation of tracks from the paired detections. We maintain a set of terminated tracks and a set of active tracks. The initial set of active tracks were obtained by adding all MB locations in the first frame as tracks of length 1. Then, we grow the tracks by processing each frame sequentially using the following procedure, for a new frame at time *t*:

- Perform MB pairing from detections in frame *t-1* to detections in frame *t* using one of the three pairing methods tested in this paper. Remove false pairings that have a pairing distance that is beyond a predefined upper limit, which was adjusted accordingly for each experiment
- For all paired MBs in frame *t*, if a detection in frame *t* is paired with a detection in frame *t-1* that exists in the set of active tracks, append the detection in frame *t* to the corresponding track.
- For all unpaired detections in frame *t*, initiate new tracks of length 1 with the detection and add them to the set of active tracks.
- For all unpaired detections in frame *t-1*, move their corresponding active tracks to the set of terminated tracks.

After processing all input frames, the set of active tracks and terminated tracks were joined together. Persistence control was applied by removing all tracks shorter than a pre-defined persistence control threshold, which was selected empirically for different experiments. To further enhance the smoothness of resulting tracks, we performed gaussian smoothing of the tracks using a 1-D gaussian filter gaussian_filter1d, which was available in the scipy.ndimage package with a sigma of 5. Vessel density maps were reconstructed by accumulating all the smoothed tracks. Velocity maps were reconstructed from using the pairing distance (after tracking smoothing) and the time interval between consecutive imaging frames. Finally, flow direction maps were determined by the angles between each pair of consecutive detections within each of the smoothed tracks, which was obtained by calculating the arc tangent of the axial displacement divided by lateral displacement between two detections. For better visibility, we presented the directional flow maps color-coded using only upward/downward directions, with upward corresponds to flow towards the transducer and downward corresponds to flow away from the transducer.

### F. Flow phantom experiment

A custom-built flow phantom was used in this study to compare the MB tracking performance among different techniques. For phantom construction, a phantom mold was first created using a clear, rectangular-shaped acrylic container with a pair of small holes drilled on opposite sides. Blunt tip dispensing needles were glued onto the outer surface of the container using epoxy resin adhesive, with the needle tubes aligned with the drilled holes. Before pouring the gelatin mixture, a 500μm diameter stainless-steel rod was inserted horizontally into the container through the dispensing tubes. The phantom was placed in a 4°C refrigerator to solidify the gelatin mixture. Prior to the imaging experiment, the metal rod was carefully removed to create a flow channel, where MB solution can be injected through one of the dispensing tubes. The diameter of the flow channel was constricted to approximately 450μm during the gelatin solidification process. DEFINITY® was diluted 1000-fold with 0.9% saline (0.9 sodium chloride, BD, Franklin Lakes, NJ) to create the MB solution perfused through the flow channel. The MB concentration in the final diluted solution is estimated to be on the order of magnitude of 10^6^MBs/mL. To generate stable MB flow with constant flow rate, one of the dispensing tubes was connected to a programmable syringe pump (NE-300, New Era Pump Systems Inc., Farmingdale, NY) with a soft plastic tube. The dispensing tube attached to the other side of the flow channel was connected to an empty beaker using a soft plastic tube. Flow rates of 20−200μL/min with 20μL increment were used in this experiment. Flow phantom data were acquired using the same transducer and imaging sequence as the PSF imaging experiment in II.C. The post-compounding frame rate was 1,000 Hz. The transducer was placed at the top surface of the phantom, positioned to provide a longitudinal view of the flow channel. 1600 frames of data (1.6s) were acquired for each flow rate.

### G. Mouse brain imaging

#### Animal model and ethics

All experimental procedures on mice were approved by the Institutional Animal Care and Use Committee (IACUC) at the University of Illinois Urbana-Champaign (protocol # 22033), and all experiments were performed in accordance with these IACUC guidelines.

#### Ultrasound imaging

Isoflurane anesthesia was induced via an induction chamber (4% isoflurane with medical oxygen) and then the mouse was transferred to a stereotaxic frame with nose cone supplying 2% isoflurane for maintenance. Lidocaine (1%) was intradermally injected under the scalp as supplementary pain relief. The mouse’s head was secured to the stereotaxic imaging frame using ear bars, and the scalp was removed with surgical scissors. Using a rotary Dremel the left side of the skull was removed starting just to the right of the sagittal suture to expose the lateral expanse of the brain.

The mouse’s tail vein was cannulated with a 30-gauge catheter and freshly activated DEFINITY® was infused at a rate of 10 µL/min using a programmable syringe pump (NE-300, New Era Pump Systems Inc., Farmingdale, NY). Ultrasound imaging was performed with the L35-16vX transducer at a center frequency of 20 MHz, using 9-angle plane wave compounding (1-degree increments) with a post-compounding frame rate of 1,000 Hz. Ultrasound data were saved as beam-formed in-phase quadrature (IQ) datasets for off-line processing in MATLAB (The MathWorks, Natick, MA; version R2019a).

### H. Rat brain imaging

#### Animal model and ethics

all animal procedures on the rat were approved by the Institutional Animal Care and Use Committee (IACUC) under protocol 22165 at the University of Illinois Urbana-Champaign.

#### Ultrasound imaging

this study used one Sprague Dawley rat (Charles River Laboratories, Inc.) of ten weeks old. The animal was subjected to isoflurane anesthesia (5% for induction and 1.5% for maintenance) throughout the experiment. After catheterizing the jugular vein, the animal was fixed on a stereotaxic frame for craniotomy. A cranial window measuring 12mm in the left-right direction and 6mm in the rostral-caudal direction was created starting from bregma. DEFINITY® microbubbles were mixed with saline to achieve a starting concentration of approximately 1.44× 10^9^ bubbles/ml. The same syringe pump used in the flow phantom experiment was used for the constant infusion of microbubble solution through the jugular vein catheter. MBs were injected with a 20µL/min flow rate. Ultrasound data were acquired using a L22-14vX transducer (Verasonics Inc., Kirkland, WA) with a 5-angle compounding (1-degree increments) plane-wave imaging sequence. The transducer was operating at a center frequency of 15.625MHz, and the post-compounding frame rate of the imaging sequence was 1,000 Hz.

## III. Results

### A. Pairing performance on simulation data

Using a simulation testing set, we quantitatively benchmarked the pairing performance of the proposed LSA pairing using fusion cost matrix (LSA-fusion) and the Bi-LSTM-based pairing against conventional LSA pairing based on minimum pairing distance (LSA-distance). Testing samples were generated using the procedure described in section II.C. Specifically, to study the behavior of pairing performance over increasing number of MBs, we generated testing samples with different MB densities ranging from of 12.5-125 MBs/mm^2^, with 100 testing samples generated for each MB density. The selection of MB concentration provides good match for the MB density observed in the *in vivo* experiments in this study. Additionally, to study the performance of each pairing algorithm under different levels of tissue motion, we simulated a set of low movement velocity of up to 98.56 mm/s and high velocity of up to 147.84 mm/s. Before combining the individual cost matrices into a fusion matrix, each individual cost matrix was scaled to a range of [0 1]. For the LSA-distance method, the weight terms *w*_*d*_, *w*_*a*_, *w*_*c*_ were set to 1, 0, 0, respectively. For the LSA-fusion method, *w*_*a*_ and *w*_*c*_ were both set to 0.2. The weights of the fusion matrix were selected empirically.

Figure 4. A and B compares the performance of LSA-distance, LSA-fusion and Bi-LSTM pairing over increasing MB concentrations for the moderate motion set and the fast motion set. Table I provides a numerical comparison of the pairing accuracy (%) at three selected concentrations of 20, 40 and 80 MBs/ mm^2^, with example sample for each concentration shown in **Figure 4**. C-E. The best pairing accuracies were highlighted in bold for each MB concentration and motion condition.

**Table 1.** Comparison of pairing accuracy (%) at select concentrations.

| Concentration | MB Velocity | LSA-distance | LSA-fusion | Bi-LSTM |
| --- | --- | --- | --- | --- |
| 20 MBs/mm<sup>2</sup> | Low | 82.7 | **87.9** | 86.1 |
|  | High | 64.6 | 74.5 | **76.6** |
| 40 MBs/mm<sup>2</sup> | Low | 66.0 | 70.1 | **72.0** |
|  | High | 44.5 | 48.9 | **54.6** |
| 80 MBs/mm<sup>2</sup> | Low | 44.5 | 46.4 | **50.1** |
|  | High | 21.8 | 23.9 | **29.4** |

Bi-LSTM achieved the best pairing performance for most scenarios, with the exception of low concentration (20 MBs/mm^2^) with low MB velocity. For low to regular MB concentrations (up to 40 MBs/mm^2^), accounting for features of MB brightness and MB shape in addition to pairwise distance in the LSA pairing framework using a simple weighted sum (i.e., LSA-fusion) greatly improves the pairing performance of LSA pairing over traditional LSA pairing with distance only. Bi-LSTM was unable to provide further notable improvement over LSA-fusion for <50 MBs/mm^2^ in the low MB movement set and <37 MBs/mm^2^ in the high movement set, as indicated by t-test. However, for higher concentrations of >60 MBs/mm^2^ in the low MB velocity set and >40 MBs/mm^2^ in the high velocity set, statistically significant improvement of Bi-LSTM pairing over LSA-fusion pairing is observed. This shows that the multi-channel Bi-LSTM pairing learns a more sophisticated combination of the provided features than simple weighted sum used by the LSA-fusion method, which enables better utilization of MB features under challenging situations of fast MB motion and high MB concentration.

**Figure 4.**
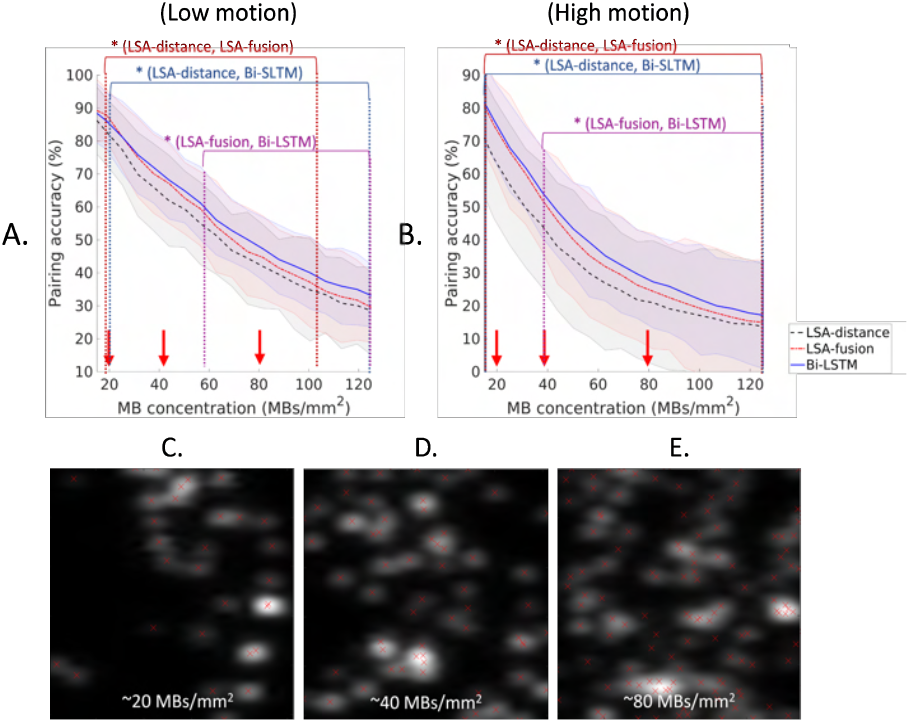
A-B. Pairing accuracy over increasing MB concentrations for different levels of motion. Shaded area indicates the standard deviation of each measurement of tracking performance for each method using the 100 testing samples at each simulated MB concentration. C-E. examples in the simulation data set at selected concentrations of 20, 40 and 80 MBs/mm^2^. The dark red, dark blue and purple dashed lines mark the intervals where the pairing accuracies are statistically significantly independent between pairing methods specified in the same colored text above. Red arrows on the x-axis in A and B correspond to the three example concentrations in C-E. Pairing accuracy at the marked concentrations are summarized in Table I.

### B. Flow phantom experiment

We then evaluated the performance of each pairing method on the gelatin flow phantom. To evaluate the accuracy of flow velocity reconstruction using each pairing method, we selected a rectangular region near the centerline of the flow channel, marked in the reconstructions in **Figure 5**. (A). The mean velocity was measured within the region for each pairing method. We excluded blank pixels in the calculation to account for the different amount of filled vessel space in each reconstruction method. The mean velocity, along with the coefficient of variation (standard deviation divided by mean) of velocity measurements within the selected region, were shown in **Figure 5**. (B). The LSA-distance velocity measurement has an obvious underestimation bias, which becomes more significant as flow velocity increases. This is expected for an algorithm that seeks to minimize total pairing distance: as MB concentration and flow velocity increases, the inter-bubble distance may become significantly smaller than the frame-to-frame displacement of a single MB, which leads to false pairing. The LSA-fusion method effectively improved upon the distance-based LSA by accounting for the MB signal shape and brightness fluctuation. Bi-LSTM pairing achieved similar improvement with slight advantage over LSA-fusion for faster flow rates of >2µL/s.

**Figure 5.**
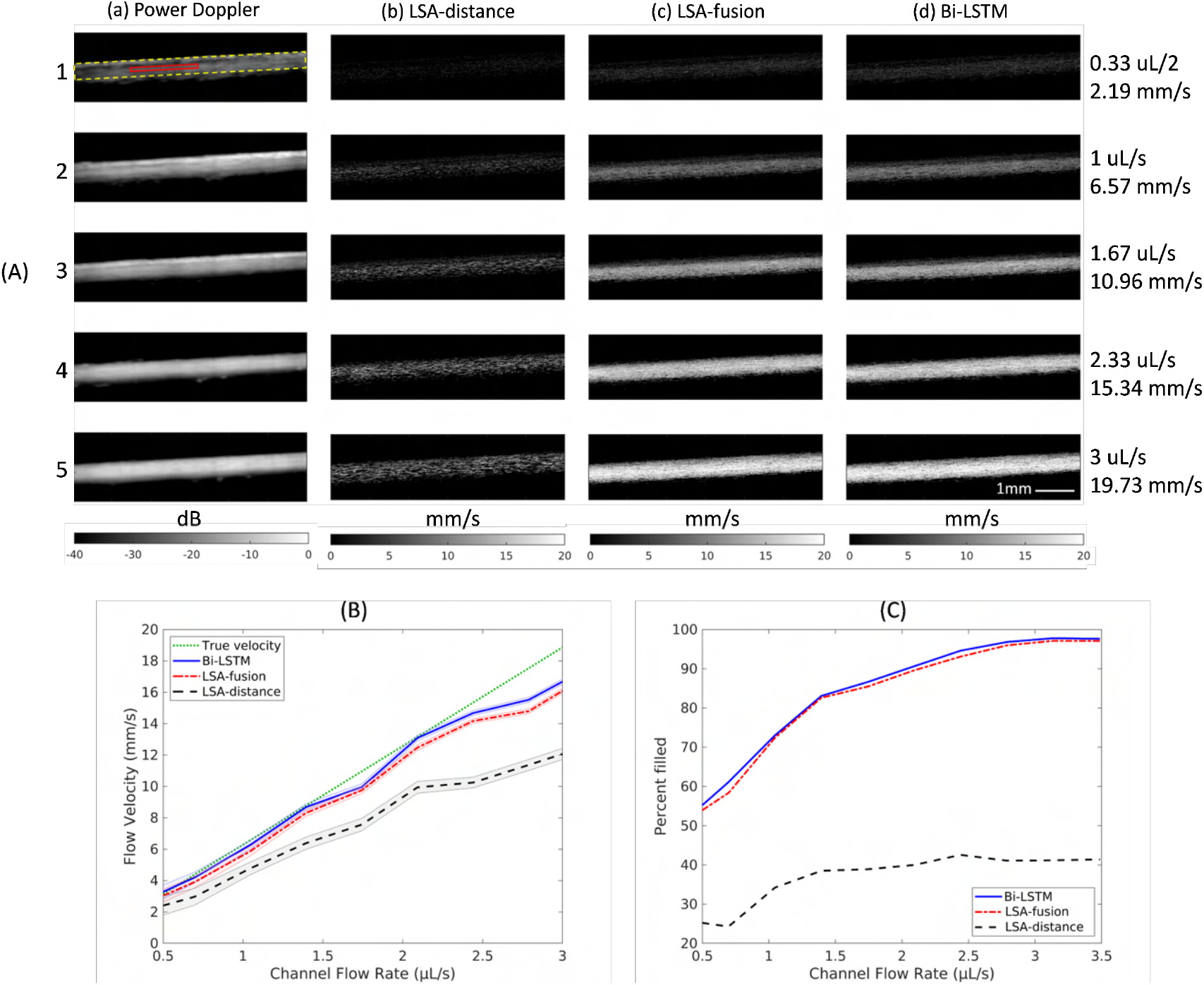
(A), Columns (a)-(d): flow phantom power Doppler images and reconstructed velocity maps based on LSA-distance, LSA-fusion and Bi-LSTM pairing of flow rates 20, 60, 100, 140 and 180 μL/s. The solid red rectangle (column (a), row 1) indicates the region for flow speed assessment. The dashed yellow rectangle was used to measure the completeness of the image reconstruction. (B) Average flow velocity measurements for each pairing method at different flow rates. The shaded area indicates the coefficient of variation for each measurement. The green dashed line indicates the ground truth velocity estimated from flow pump. (C) Percentage of pixels covered by MB tracks in the final reconstructed velocity map.

In addition to the accuracy of flow velocity estimation, we also investigated each pairing method’s efficacy at producing reliable MB tracks that effectively map out the perfused vascular space. We evaluated this criterion by calculating the percentage of pixels filled by MB tracks in the final reconstructed velocity map. **Figure 5**. (C) shows the results. Because small pairwise distance is preferred by LSA-distance pairing, a large amount of incorrectly linked tracks that jitter in a small local region were produced, which were then later discarded by persistence control (10ms for the flow phantom experiment). As a result, many detected MBs were not able to contribute to the reconstruction of the final velocity map, resulting in very sparse reconstruction across all flow rates. On the other hand, both LSA-fusion and Bi-LSTM were able to generate long MB tracks with more robust pairing, leading to significantly improved filling rates. This result indicates that the additional information of MB brightness and shape enabled more consistent and accurate MB tracking, which facilitated more robust reconstruction of the final velocity map. Interestingly, Bi-LSTM did not demonstrate significantly better performance than LSA-fusion in the flow phantom study. This result may be attributed to the fact that the flow phantom contains a rather simple flow pattern where MBs move consistently towards the same direction, which is significantly less challenging than *in vivo* blood vessels. While both LSA-fusion and Bi-LSTM were able to achieve similar level of improvement over LSA-distance by using additional information contained in the MB signal, the advantage of Bi-LSTM will further manifest *in vivo*.

### C. In vivo mouse brain experiment

We first validated the proposed tracking algorithms *in vivo* in the mouse brain. Different pairing algorithms were applied to the same set of MB localization data obtained using the method described in II.E. To reject erroneous tracking results, we applied a threshold of 10 pixels on the maximum MB movement between consecutive frames, which is equivalent to a maximum MB velocity of 49.28mm/s. The persistence control was set to 10ms for all methods. **Figure** 6 (a)-(c) shows the reconstructed velocity maps based on LSA-distance pairing, LSA-fusion pairing, and Bi-LSTM pairing. Overall, the LSA-distance pairing resulted in velocity map reconstructions with lowest quality in the context of vessel completeness and variations in flow velocity estimation. This observation matches with the observations in the flow phantom experiment.

For the cortical region (Error! Reference source not found.**Figure 6**(a) Box1), closely spaced small cortical vessels pose the most significant challenges for tracking: MB events can be easily associated with wrong localizations in nearby vessels, resulting in noisy reconstruction and isolated and fragmented MB tracks (e.g., Figure 6(d) and (g)). Bi-LSTM reconstruction (**Figure** 6 (f) and (i)), on the other hand, demonstrated a better pairing performance as evidenced by the reduced number of fragmented tracks and the smoothest appearance of vessels. This improvement may be attributed to the more efficient and accurate use of MB localizations towards the reconstruction of vessel maps (as opposed to false pairings that result in tracks with low persistence that were later discarded by persistence control). We selected two pairs of vessels (1 and 2 in **Figure** 6 (d)) to further examine the velocity profile (**Figure** 6 (p) and (q)). **Figure** 6 (p) shows two closely spaced vessels with opposite flow direction and similar flow velocity. LSA-distance pairing was able to reconstruct the positive flow with clear evidence of a parabolic flow profile.

**Figure 6.**
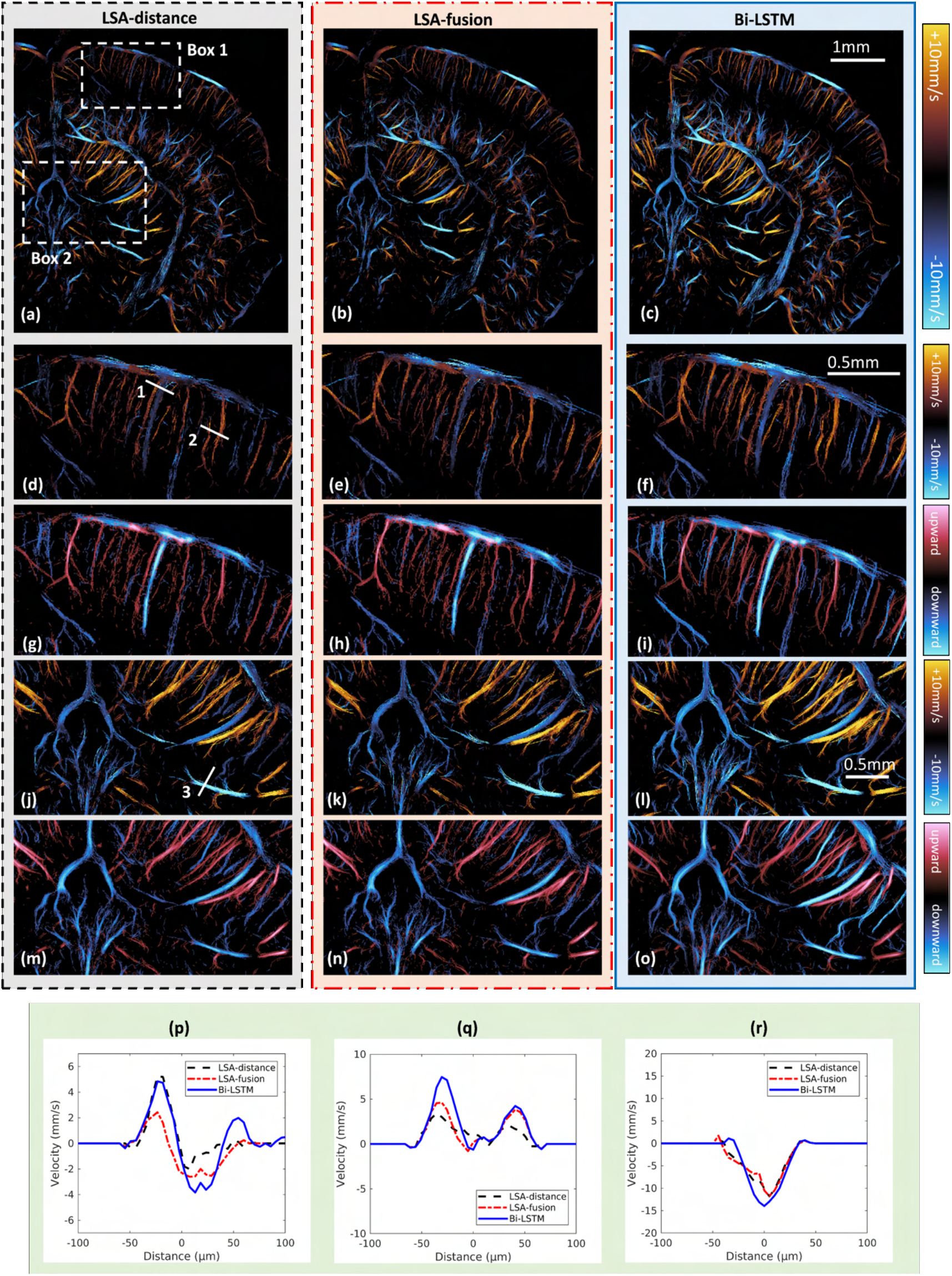
(a)-(c) Super-resolved velocity maps reconstructed based on LSA-distance pairing, LSA-fusion pairing and Bi-LSTM pairing using 102.4s of contrast enhanced mouse brain data. Box 1 and Box 2 are ROIs selected for further zoom-in examination. (d)-(f) zoom in directional flow of Box 1 in (a) reconstructed using LSA-distance, LSA-fusion, and Bi-LSTM pairing. (g)-(i) are the corresponding directional velocity maps. (j)-(l) directional flow of Box 2 reconstructed using LSA-distance, LSA-fusion, and Bi-LSTM pairing. (m)-(o) are the corresponding directional velocity maps. (p)-(r) Comparison of flow velocity profiles reconstructed using LSA-distance pairing, LSA-fusion pairing, and Bi-LSTM pairing of profiles 1-3 marked in (d) and (j).

However, it produced a suboptimal reconstruction of the vessel with opposite flow direction. LSA-fusion, although being able to clearly distinguish the two vessels with opposite flow directions, reconstructed both vessels with lower flow velocity. Bi-LSTM was able to produce a clean separation of both vessels with clear parabolic flow profiles. Based on the observation in the flow phantom experiment, we believe it is more likely that the LSA-fusion underestimated the flow velocity due to false tracking. **Figure** 6 (q) corresponds to two closely spaced small vessels with the same flow direction and slightly different flow velocity. Again, both LSA-fusion and Bi-LSTM were able to better separate the two vessels compared to LSA-distance. Similar to the case in **Figure** 6 (p), LSA-fusion underestimated the velocity of the faster vessel. Bi-LSTM provided the best separation with clear parabolic flow profiles for both vessels.

The thalamic region (**Figure** 6 (a) Box 2) includes bigger blood vessels with faster flow and higher MB concentrations, which presents another challenging for MB tracking. In addition, due to it being further from the transducer, the thalamic region also has lower MB SNR. Due to the abundance of MB events in this region, all three methods were able to reconstruct most of the closely spaced vessels in the 10-50 µm range. However, both LSA-based methods were susceptible to generating faulty tracks under the influence of noise, resulting in a fuzzier appearance to the reconstructed vessel maps. In contrast, Bi-LSTM was able to produce the cleanest and smoothest reconstruction of all types of vessels in this region. Additionally, as seen in the directional flow map, Bi-LSTM produced the least amount of track fragments around the clearly vascularized regions, exhibiting superior robustness to noise. **Figure** 6 (r) shows the comparison of reconstructed vessel profiles of the vessel labeled 3 in **Figure** 6 (j). Similar as the observations above, Bi-LSTM was able to recover a clear parabolic flow profile for the targeted vessel.

### D. In vivo rat brain experiment

Finally, we validated the proposed tracking algorithms on a set of *in vivo* rat brain data. The rat brain data was acquired using a different transducer (L22-14vX, Verasonics, Kirkland, WA) with a center frequency of 15 MHz, which is different from the simulation data used for network training. Therefore, this experiment provides a test for the generalizability for the proposed deep learning-based MB tracking method.

The persistence control for MB tracking was set to 20 ms for LSA-fusion and Bi-LSTM pairing. For LSA-distance pairing, we noticed that a persistence control of 20ms rejected the majority of the tracks, resulting in an extremely fragmented reconstruction. Therefore, we elected to set a more relaxed persistence control of 10ms for LSA-distance pairing. **Figure 7**(a)-(c) shows the reconstructed velocity maps and **Figure 7**(d)-(f) shows the directional flow maps. LSA-distance pairing resulted in the sparsest velocity map reconstruction even with the most relaxed persistence control, which agrees with the observations in the phantom and mouse brain imaging experiments. For both the velocity map and the directional vessel map, Bi-LSTM delivered the most complete reconstruction with clear delineation of individual vessels.

**Figure 7.**
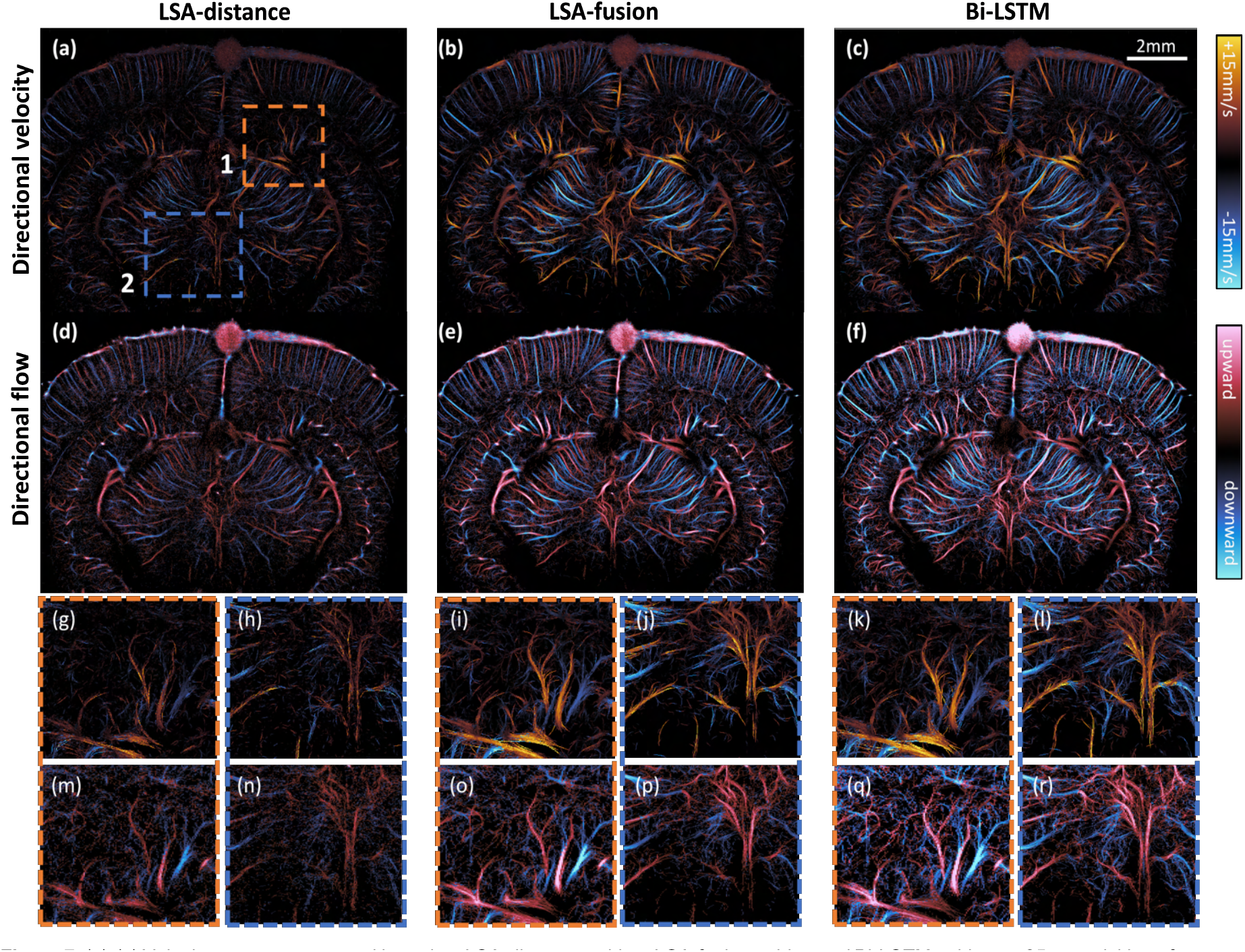
(a)-(c) Velocity map reconstructed based on LSA-distance pairing, LSA-fusion pairing and Bi-LSTM pairing on 25s acquisition of contrast enhanced rat brain data. (d)-(f) directional flow map reconstructed based on LSA-distance pairing, LSA-fusion pairing and Bi-LSTM pairing on 25s acquisition of contrast enhanced rat brain data. (g)-(l), enlarged version of the velocity maps of regions 1 and 2 in (a). (m)-(r), enlarged version of the directional flow maps of regions 1 and 2 in (a).

For region 1 labeled in **Figure 7**(a), MB events were frequently captured within vessels with under 50µm diameter in between the major branches. Due to the sporadic nature of such events, LSA-based methods were unable to consistently track the movement of a single MB for a long time, resulting in insufficient number of tracks that can be used in the final reconstruction. The directional flow map based on Bi-LSTM pairing (**Figure 7**(q)) shows clear tracings of MB movement trajectories within the small vessels, indicating improved tracking robustness. In the deeper thalamic region (region 2 in **Figure 7**(a)), the raw MB data had the lowest SNR, which results in noisy and incomplete reconstructions by both LSA-based methods. Similar to the mouse brain study, Bi-LSTM exhibited improved robustness to noise and was able to successfully reconstruct the deeper region with clear vascular structures.

Results presented in **Figure 7** also indicate that the proposed Bi-LSTM method has robust generalizability for different imaging settings on different animal models. The robust adaptability was anticipated because the Bi-LSTM model learned to predict from the relationship between each MB pairs rather than the specific features of the MB signal (i.e., the input to the network is not MB signal as shown in **Figure 1**). Therefore, although trained using simulation data generated from a particular imaging setup, the performance of the proposed Bi-LSTM pairing method did not drop when applied to data acquired from a different experimental setting. Once trained, the proposed method should be easily adapted to different datasets with minimum tuning.

## IV. Discussion

In this paper, we proposed a deep learning-based MB pairing method for MB tracking using the Bi-LSTM neural network. The proposed method made efficient use of information contained in the ultrasound MB data to produce more reliable pairing to enhance ULM reconstruction. The proposed method outperformed LSA-based tracking based on MB pair-wise distance, which is commonly used method in the ULM community. It also outperformed LSA-based tracking using a fusion cost matrix, which utilizes more information (e.g., MB signal amplitude and shape) in addition to distance to assist pairing. Additionally, Bi-LSTM-based pairing does not require manual tuning of feature weights of LSA-based tracking with fusion cost matrix (Equation (3)), which avoids operator-induced biases.

In our deep learning-based pairing formulation, the functionality of the Bi-LSTM module was somewhat atypical. Specifically, the input sequence to the Bi-LSTM layers was not a temporal sequence in the regular sense, but rather the sequence of the possible association costs for a MB detection in one frame. Instead of extracting long-term temporal features, the Bi-LSTM layers act as feature extraction modules with large receptive field, through memorizing information associated with previous inputs in the cell state in the forward LSTM, and subsequent inputs in the cell state in the backward LSTM. It was discussed in the design of Deep Hungarian Network [35] that CNNs and regular RNNs can also be used to fulfill the basic requirements of a DL-based formulation of the bi-partite pairing problem. We believe that Bi-LSTMs are better suited for this task: they do not have the inherent limitation of receptive fields of CNNs and are able to better retain long-term memory compared to regular RNNs. However, it should be noted that the ability of retaining information in the LSTM is limited by the size of its internal states (cell states and hidden states). Despite theoretically being able to handle arbitrary length input, the hidden dimensions of LSTM may need to be adjusted accordingly when processing inputs where the number of detections of each input frame is very large. Additionally, for the two-frame pairing formulation, the order within a sequence of assignment cost that is being fed into the Bi-LSTM layers is not very relevant to the result. Thus, some of the internal memory of the LSTM may be used to retain irrelevant information. Ideally, the model would learn to ‘forget’ such redundant information through its gated mechanism. Additional regularization can be introduced to the training process to alleviate this issue. The Bi-LSTM layer can potentially be replaced by a transformer [38], but memory consumption may become a concern due to their high memory complexity.

The proposed Bi-LSTM pairing has practical limitations. For computing time, our current implementation of Bi-LSTM pairing requires roughly two times the computational time compared to LSA-based pairing, which can become a significant hurdle for large-scale processing. Additionally, integration of a DL-based pairing into an existing ULM processing pipeline may involve challenges associated with data transfer, cross platform performance, and miscellaneous overheads. Therefore, we believe that the LSA-fusion pairing method can be a good substitute to quickly boost the tracking performance of an existing ULM processing chain with minimum developmental overhead.

The current formulation of the proposed method only considers association of MBs between two consecutive frames *t* and *t* + 1. It has the potential to be extended to a multi-frame association method by expanding the NN input and output to contain association cost and assignment matrix of multiple frames (*t* to *t* + 1, t to *t* + 2, etc.). However, the track formation from multi-frame assignment will need to be modified and will introduce additional computation complexity.

The performance of Bi-LSTM pairing method is largely affected by the quality of the training data. At present, the pipeline for generating simulation data to train the Bi-LSTM model aims to include most generic cases for better generalizability. Therefore, we did not assume any motion model, nor did we apply any underlying vessel structure to the spatial distribution of the MBs. Generating high-quality training data with tissue specific structure and flow dynamics for fine-tuning or continued training of a generic model can potentially further improve the performance for specific type of tissue. Additionally, the current simulation pipeline did not enforce a match between the spatial location of the PSFs in the experimental data and where they were used in the simulation data. An experimental PSF from a certain depth could be used to generate MB image at a different depth. Due to the high framerate of our acquisition, the velocity range of our in vivo data, and the displacement thresholding on the paired MBs, we decided that the spatial variation of PSF will not significantly affect the pairing process. If the proposed method is to be applied to lower framerate data or much faster flow, the simulation setup may need to be redesigned to consider the spatial variability of PSFs.

The Bi-LSTM tracking component can potentially be used in conjunction with a DL-based localization component for joint training and an end-to-end DL-based implementation of ULM. However, chaining DL models may result in training complications due to issues associated with vanishing or exploding gradient. The training procedure will need to be carefully designed. Additionally, combining two DL modules may amplify their corresponding limitations. If used in combination with a DL-based localization method, the performance of the combined ULM pipeline will need to be carefully evaluated against existing benchmark methods.

## V. Conclusion

We proposed a novel microbubble (MB) pairing method for super-resolution ultrasound localization microscopy (ULM) based on a Bi-LSTM neural network. The proposed method achieved more robust and accurate MB pairing performance by integrating MB signal features into the pairing process. It also minimizes operator biases in designing the pairing criteria. The proposed method demonstrated the best MB pairing and tracking performance in the *in silico, in vitro*, and *in vivo* experiments presented in this study with varying MB concentrations and flow conditions.

